# Adipocyte-secreted IL-6 sensitizes macrophages to IL-4 signaling

**DOI:** 10.1101/2022.07.19.500620

**Authors:** Danny Luan, Benyamin Dadpey, Jessica Zaid, Pania E. Bridge-Comer, Julia H. DeLuca, Wenmin Xia, Joshua Castle, Shannon M. Reilly

**Author notes:** Corresponding author: Shannon M. Reilly. Trutino Biosciences, San Diego, CA 92121, USA. Virginia Tech Carilion School of Medicine, Roanoke, VA, USA. Department of Orthopaedic Surgery, Henry Ford Hospital, Detroit, Michigan, U.S.A. These authors contributed equally (Danny Luan, Benyamin Dadpey).

## Abstract

Complex bidirectional crosstalk between adipocytes and adipose tissue immune cells plays an important role in regulating adipose function, inflammation, and insulin responsiveness. Adipocytes secrete the pleiotropic cytokine IL-6 in response to both inflammatory and catabolic stimuli. Previous studies suggest that IL-6 secretion from adipocytes in obesity may promote adipose tissue inflammation. Here we investigated catabolic stimulation of adipocyte IL-6 secretion and its impact on adipose tissue immune cells. In obesity, catecholamine resistance reduces cAMP-driven adipocyte IL-6 secretion in response to catabolic signals. By restoring adipocyte catecholamine sensitivity in obese adipocytes, amlexanox stimulates adipocyte-specific IL-6 secretion. Here we report that in this context, adipocyte secreted IL-6 activates local macrophage STAT3 to promote *Il4ra* expression, thereby sensitizing them to IL-4 signaling, and promoting an anti-inflammatory gene expression pattern. Supporting a paracrine adipocyte to macrophage mechanism, these effects could be recapitulated using adipocyte conditioned media to pretreat bone marrow derived macrophages prior to polarization with IL-4. The effects of IL-6 signaling in the adipose tissue are complex and context specific. These results suggest that cAMP driven IL-6 secretion from adipocytes sensitizes adipose tissue macrophages to IL-4 signaling.

---

Chronic low-grade inflammation has been implicated in many of the common co-morbidities associated with obesity, including type 2 diabetes and non-alcoholic steatohepatitis. The non-canonical I-κB kinases IKKε and TBK1 are induced by chronic inflammation in adipocytes and perpetuate obesity by affecting catecholamine resistance via the activation of phosphodiesterase-3B [1-4]. Treatment of obese mice with amlexanox, a dual specific inhibitor of IKKε and TBK1, results in improved metabolic health via weight loss and resolution of white adipose tissue (WAT) inflammation [2]. By restoring cAMP signaling in adipocytes, amlexanox also promotes the expression of IL-6 [5]. IL-6 is a pleotropic cytokine with a complex role in obesity and WAT inflammation. While IL-6 levels are associated with increased risk of diabetes in obese individuals [6-8], *Il6* knockout (KO) mice are protected from age-associated obesity and diet-induced metabolic dysfunction [9, 10]. And IL-6 signaling in macrophages has an anti-inflammatory impact in obese WAT [11, 12]. The specific contribution of adipocyte-secreted IL-6 is unclear: one study observed proinflammatory trans-signaling in obese WAT [13], while another observed no effect [14]. Catabolically stimulated adipocyte IL-6 secretion has not been investigated.

In this study, we found that in vivo amlexanox treatment resulted in an IL-6 dependent Tyr705 phosphorylation of STAT3 in adipose tissue macrophages (ATMs). The activation of STAT3 in macrophages resulted in upregulation of *Il4ra*, and sensitization to IL-4 signaling. This effect could be recapitulated in vitro by treating bone marrow-derived macrophages (BMDMs) with conditioned media from adipocytes treated with amlexanox. In vivo amlexanox treatment sensitized ATMs to IL-4 in a macrophage STAT3 dependent manner.

## Research Design and Methods

### Animals

The following strains of mice from Jackson Labs were bred for littermate-controlled experiments: C57BL/6J (RRID: IMSR_JAX:000664) *Stat3* floxed (RRID: IMSR_JAX:016923), *LysM-cre* (RRID: IMSR_JAX:004781), *Adipoq-cre* (RRID: IMSR_JAX:010803), *Il6 KO* (RRID: IMSR_JAX:002650). Obesity was induced by high fat diet (HFD), 45% of calories from fat (D12451, Research Diets), starting at 6-10 weeks of age. Amlexanox was administered by daily oral gavage at a dose of 25□mg/kg to preconditioned male mice after 13-14 weeks of HFD. Mice were housed in a specific pathogen-free facility with a 12-h light, 12-h dark cycle, and given free access to food and water. All animal use was approved by the Institutional Animal Care and Use Committee (IACUC) at the University of California-San Diego, University of Michigan, and Weill Cornell Medicine.

Stromal vascular cells (SVC) and mature adipocytes were isolated from WAT by centrifugation following collagenase digestion. Serum IL-6 levels were quantified using Mouse IL-6 Quantikine ELISA from R&D (SM6000B) with 50 μl of serum or media.

### Cells

#### 3T3-L1 media conditioning

RPMI media from differentiated 3T3-L1s treated with 100 μM amlexanox or vehicle was collected after 4 hours. Conditioned media were incubated with IL-6 NA 10 μg/mL (R&D Systems Cat# MAB406, RRID:AB_2233899) or normal goat IgG control antibody 10 μg/mL (R&D Systems Cat# AB-108-C, RRID:AB_354267) for 15 min.

#### BMDM

Dispersed bone marrow cells from 6-8 week old male mice were placed in culture media (RPMI 1640, 10% FBS, MCSF 20 ng/mL (Peprotech, no. 315-02), 20 mM HEPES, 2 mM L-Glutamine, and 1 mM Sodium Pyruvate). On day 6, cells were incubated with 50 ng/mL IL-6 (R&D, no. 406-ML-025) or culture media made with conditioned RPMI prior to the addition of 10 ng/mL IL-4 (R&D, no. 404-ML-025/CF) on day 7.

### Fluorescence activated cell sorting (FACS)

WAT macrophages identified as CD45^+^, CD11c^+^, Emr1^+^, CD31^-^ were sorted into triazole for gene expression analysis. Each animal was sorted separately. BMDM identity was confirmed by Emr1 and CD11b dual positivity. The following antibodies were purchased from Thermo Fisher Scientific: FC block (CD16/32) (14-0161-82, RRID:AB_467133), FITC conjugated anti-CD45(30-F11) (11-0451-82, RRID:AB_465050), Pacific Blue conjugated anti-CD45(30-F11) (MCD4528, RRID:AB_10373710), Brilliant Violet conjugated anti-CD45 (103147, RRID:AB_2564383), PE conjugated anti-CD64 (12-0641-82, RRID:AB_2735014), APC conjugated anti-CD11c (17-0114-82, RRID:AB_469346), Percp-Cy5 conjugated anti-CD3 (45-0031-82, RRID:AB_1107000), PE-Cy7 conjugated anti-CD31 (25-0311-82, RRID:AB_2716949), APC-Cy7 conjugated anti-LysG (25-9668-82, RRID:AB_2811793), eFluor-450 conjugated pTyr705 STAT3 (48-9033-42, RRID:AB_2574121). The following antibodies were purchased from BioLegend: APC-Cy7 conjugated anti-CD11c (117324, RRID:AB_830649), APC conjugated anti-Emr1 (123116, RRID:AB_893481), APC/Fire conjugated anti-CD11b (101262, RRID:AB_2572122), and PE/Cy7 conjugated anti-CD31 (102418, RRID:AB_830757). Alexa Fluor 647 conjugated anti-CLEC10A/CD301, from Novus Biologicals, (64874AF647, RRID: 2915978). Stained cells were analyzed or sorted by flow cytometry on a MoFlo Astrios cell sorter (Beckman Coulter) at the University of Michigan’s Flow Cytometry Core, or on the BD FACS Aria II at UCSD’s Flow Cytometry Core in Moores Cancer center.

### Western blot analysis

Primary antibodies were purchased from Cell Signaling Technology: pTyr-705 STAT3 1:1000 (9145, RRID:AB_2491009), STAT3 1:4000 (9139, RRID:AB_331757) STAT6 (9362, RRID:AB_2271211), pTyr-641 STAT6 1:1000 (56554, RRID:AB_2799514) and *β*-Tubulin 1:1000 (2146, RRID:AB_2210545), p38 1:1000 (9212, RRID:AB_330713). Secondary antibodies were purchased from Thermo Fisher Scientific: Goat anti-mouse 1:10,000 (31430, RRID:AB_228307) and goat anti-rabbit 1:10,000 (31460, RRID:AB_228341).

### Immunohistochemistry

After blocking with 5% goat serum, slides were incubated overnight with 1:200 anti-pTyr705 STAT3 and detected with DAB after washing.

### Gene expression Analysis

Real-time PCR with Power SYBR Green was performed using the Applied Biosystems QuantStudio5 real-time PCR System and quantified using an internal standard curve with *Arbp* as the control gene.

### Data and Resource Availability

All data generated or analyzed during this study are included in the published article (and its online supplementary files). No applicable resources were generated or analyzed during the current study.

## Results

Amlexanox treatment of obese mice acutely promotes adipocyte IL-6 secretion via cAMP signaling [15]. Four hours after oral gavage with amlexanox, when serum IL-6 levels peak, tyrosine 705 phosphorylation (pTyr705) of STAT3 is elevated in WAT (Fig. 1A). Immunohistochemical analysis revealed many pTyr705 STAT3 positive nuclei in crownlike structures (Fig. 1B). KO of STAT3 in adipocytes significantly reduced total WAT levels of STAT3 (Fig. 1A and C). The remaining STAT3, attributable to non-adipocyte cells, was similarly phosphorylated (Fig. 1A and D). To investigate STAT3 phosphorylation in immune cells, SVCs were collected and analyzed for pTyr705 STAT3 by fluorescence activated cell sorting.. Amlexanox specifically increased pTyr705 STAT3 in both Cd11c^+^ and Cd11c^-^ adipose tissue macrophages (ATMs) (Fig. 1E-G). No STAT3 activation was observed in neutrophils, dendritic cells or T cells (Fig. 1H-J). The mechanism of specificity for macrophages is not clear.

**Fig. 1.**
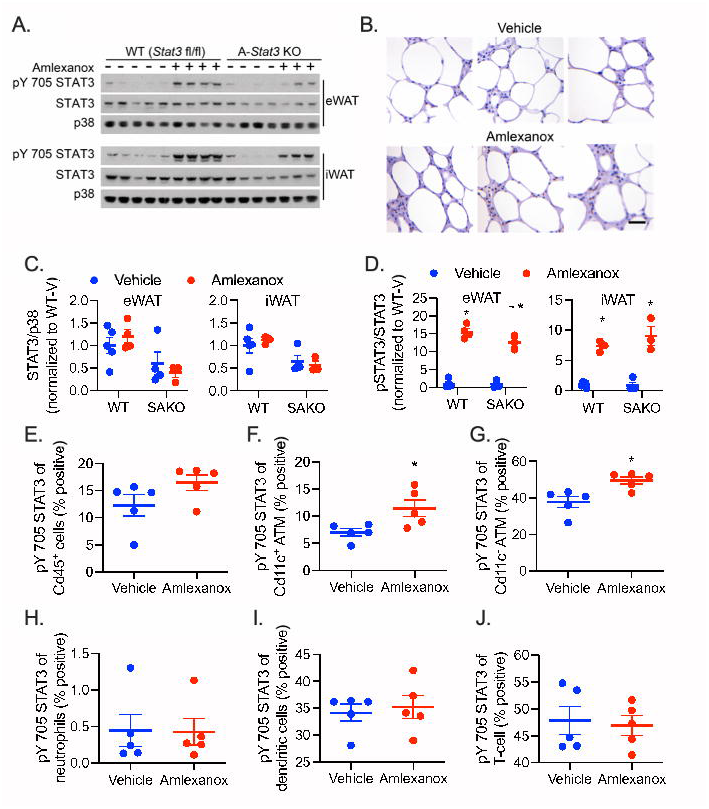
STAT3 phosphorylation in adipose cells after amlexanox treatment. Experiments were performed 4 h after gavage with 25 mg/kg amlexanox or vehicle control in obese male mice aged 20-24 weeks. (A) Western blot analysis of STAT3 and Tyr705 phosphorylated STAT3 (pSTAT3) in epididymal and inguinal white adipose tissue (eWAT and iWAT respectively of adipocyte specific KO mice and floxed littermate controls. (B) Immunohistochemical analysis of pTyr705 STAT3, brown DAB staining. Slides also stained with H&E. Tissues harvested and immediately fixed after 52 h amlexanox treatment by daily oral gavage. Scale bar = 50 *μ*m. n = 3 per treatment. (C) Quantification of STAT3 over p38 levels in eWAT (left) and iWAT (right). The effect of genotype is significant in both tissues, p < 0.01 by two-way ANOVA. (D) Quantification of pY705 STAT3 over total STAT3 levels in eWAT (left) and iWAT (right). * p < 0.05 vehicle versus amlexanox within genotype. ^∼^ p < 0.05 WT versus KO within treatment group. (E-J) FACS analysis of the percent positivity for pTyr705 STAT3 in SVC populations. N = 5 per treatment. * p < 0.05 by student’s t-test vehicle versus amlexanox. (E) All Cd45 positive immune cells. (F) Proinflammatory ATMs = Cd45^+^, Cd64^+^, Cd11c^High^. (G) Anti-inflammatory ATMs = Cd45^+^, Cd64^+^, Cd11c^Low^. (H) Neutrophils = Cd45^+^, Ly6G^+^. (I) Dendritic cells = Cd45^+^, Cd64^-^, Cd11c^+^. (J) T-cells = Cd45^+^, Cd3^+^. Statistical significance determined by post hoc analysis after significant two-way ANOVA.

To determine dependence on IL-6, we utilized whole body *Il6* KO animals. While littermate controls demonstrate elevated serum IL-6 following amlexanox treatment, IL-6 levels were undetectable in the *Il6* KO animals (Fig. 2A). Hepatic STAT3 is activated by adipocyte-secreted IL-6 following amlexanox treatment [15]. As expected, amlexanox treatment induced hepatic pTyr705 STAT3 in the WT but not *Il6* KO livers (Fig. 2B and C). Staining for pTyr705 STAT3 was prominent in crown-like structures from WT but not *Il6* KO WAT (Fig. 2D and E). Accordingly, *Socs3*, a STAT3 target gene, was elevated in the WAT by amlexanox treatment only in WT mice (Fig. 2F). The increase in macrophage pTyr705 STAT3 by amlexanox was dependent on IL-6, as it was not observed in ATMs from *Il6* KO mice (Fig. 2G-K). The presence of pTyr705 STAT3 in the *Il6* KO ATMs may be mediated by another IL-6 family cytokine, none of which are induced by amlexanox (Supplementary Fig. 1).

**Fig. 2.**
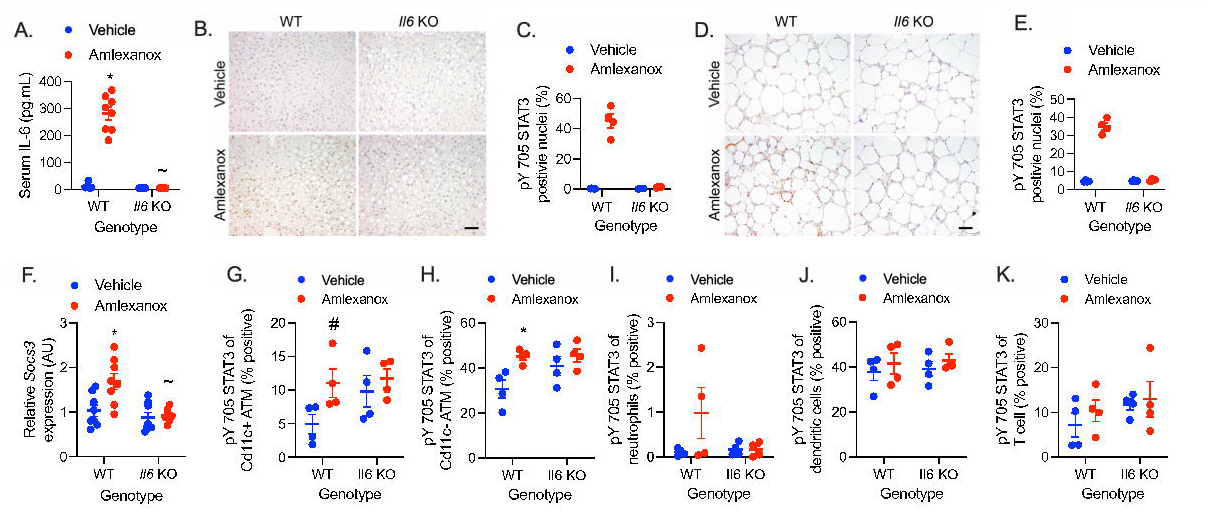
STAT3 activation in adipose tissue immune cells is IL-6 dependent. Experiments were performed 4 h after gavage with 25 mg/kg amlexanox or vehicle control in obese *Il6* KO and WT littermate control male mice aged 20-24 weeks. (A) Serum IL-6 levels. n = 8 per group. (B-E) Immunohistochemical analysis of pTyr705 STAT3, brown DAB staining. Slides also stained with Hematoxylin. Tissues harvested and immediately fixed 4 hours after gavage. (B-C) Liver. (D-E) WAT. (B and D) Representative image from each genotype and treatment, scale bar = 100 *μ*m. (C and E) Quantification of percent positive nuclei in sections from four animals per condition (three fields of view with approximately 200 nuclei each were averaged for each animal, n = 4 per group). (F) Relative expression of *Socs3* in WAT. n = 8 per group. (G-K) FACS analysis of the percent positivity of pTyr705 STAT3 in SVC populations. n = 4 per group. (G) Proinflammatory ATMs = Cd45^+^, Cd64^+^, Cd11c^High^. (H) Anti-inflammatory ATMs = Cd45^+^, Cd64^+^, Cd11c^Low^. (I) Neutrophils = Cd45^+^, Ly6G^+^. (J) Dendritic cells = Cd45^+^, Cd64^-^, Cd11c^+^. (K) T-cells = Cd45^+^, Cd3^+^. Statistical significance determined by post hoc analysis after significant two-way ANOVA. * p < 0.05 vehicle versus amlexanox within genotype. ^∼^ p < 0.05 WT versus KO within treatment group. ^#^ p < 0.05 vehicle versus amlexanox within genotype using Fisher’s exact test.

IL-6 stimulation in macrophages activates STAT3-mediated expression of *Il4ra*, thereby sensitizing the cells to anti-inflammatory effects of IL-4 [11, 12]. Using bone marrow-derived macrophages (BMDM), we confirmed that IL-6 treatment increased *Il4ra* expression and pTyr-641 STAT6 in response to IL-4 treatment (Supplementary Fig. 2A and B). IL-6 pretreatment increased the induction of *Arg1* and suppression of *Il1b* expression by IL-4 (Supplementary Fig. 2C and D).

To determine whether IL-6 secreted from amlexanox-treated adipocytes may function as a paracrine signal to ATMs, we treated BMDM with conditioned media from amlexanox-treated 3T3-L1 adipocytes (Fig. 3A). Amlexanox-conditioned media (ACM), but not direct amlexanox treatment, increased the percentage of CD301^+^ cells in BMDM polarized with IL-4 (Fig. 3B). While conditioned media from adipocytes treated with vehicle control (VCM) did not significantly increase *Il4ra* expression in BMDM, conditioned media from adipocytes treated with amlexanox (ACM) increased *Il4ra* expression six-fold over non-conditioned control media (NCM) (Fig. 3C). ACM but not VCM increased *Arg1* and decreased *Itgax* expression in BMDM polarized with IL-4 (Fig. 3D and E). Importantly, ACM did not promote CD301^+^ macrophages in *Stat3* KO BMDM (from LysM-Cre driven myeloid cell-specific *Stat3* KO (SMKO) mice), indicating that the effects of the ACM on macrophage polarization were mediated through STAT3 (Fig. 3B). Direct amlexanox treatment did not promote *Il4ra* or *Arg1* expression in BMDMs, either alone or in combination with IL-6 treatment (Fig. 3F and G). While *Itgax* expression was additively suppressed both by IL-6 and amlexanox (Fig. 3H).

**Fig. 3.**
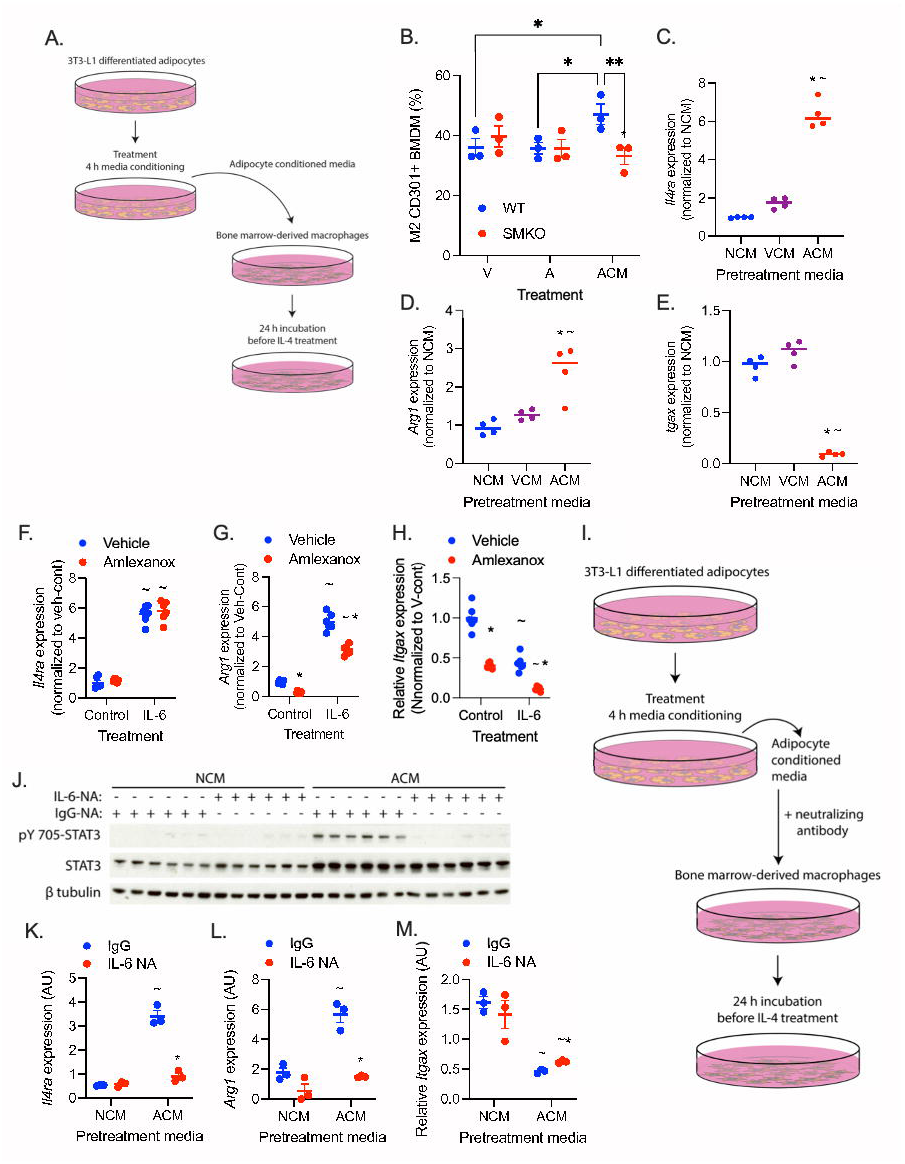
Adipocyte-secreted IL-6 sensitizes macrophages to IL-4. Adipocyte-conditioned media was generated by treating adipocytes with 100 μM amlexanox in RPMI for 4 hours. Direct treatment of BMDMs with amlexanox was also performed with 100 μM amlexanox. (A) Schematic of adipocyte media conditioning and treatment of BMDMs. (B) Percent CD301^+^ staining of F4/80, CD11b dual positive BMDM treated with amlexanox directly, or conditioned media from amlexanox treated adipocytes (ACM). n = 3 per group. * p < 0.05 comparison indicated by line. (C-E) Gene expression in BMDMs pretreated with non-conditioned media (NCM), vehicle conditioned media (VCM) or amlexanox-conditioned media (ACM) for 24 h before the addition of IL-4 for another 24 h. n = 4 per group. * p < 0.05 ACM versus VCM, ^∼^ p < 0.05 ACM versus NCM (C) Relative *Il4ra* expression. (D) Relative *Arg1* expression. (E) Relative *Itgax* expression. (F-H) Gene expression in BMDMs treated with 50 ng/mL IL-6 with and without amlexanox, normalized to the vehicle control untreated condition. n = 6 per group. * p < 0.05 vehicle versus amlexanox, ^∼^ p < 0.05 control versus IL-6. (F) Relative *Il4ra* expression. (F) Relative *Arg1* expression. (F) Relative *Itgax* expression. (I) Schematic of adipocyte media conditioning with neutralizing antibodies and administration to BMDMs. (J-M) BMDM treated with control or amlexanox conditioned media in which IL-6 was neutralized with IL-6NA or IgG control. * p < 0.05 IgG versus IL-6NA, ^∼^ p < 0.05 ACM versus NCM. (J) Western blot analysis of STAT3 Tyr705 phosphorylation, *β*-Tubulin serves as a loading control. (K) Relative *Il4ra* expression. (L) Relative *Arg1* expression. (M) Relative *Itgax* expression. n = 3 per group. Statistical significance determined by post hoc analysis after significant ANOVA.

To confirm that IL-6 in the ACM mediates the effects of amlexanox, we treated the conditioned media with IL-6 neutralizing antibodies (IL-6NA), or IgG antibody control (Fig. 3I and Supplementary Fig. 3). Neutralization of IL-6 prevented the induction of pTyr705 STAT3 and blocked the induction of *Il4ra* in BMDM treated with ACM (Fig. 3J and K). Accordingly, neutralization of IL-6 in the ACM blocked its ability to promote *Arg1* expression in IL-4 polarized macrophages and attenuated the suppression of *Itgax* expression (Fig. 3L and M). These results suggest that IL-6 secreted from amlexanox-stimulated adipocytes sensitizes macrophages to IL-4 signaling.

To determine if in vivo amlexanox treatment results in similar IL-4 sensitization in ATMs, we measured gene expression in mature adipocytes and ATMs isolated from mice 8 hours after a single dose of amlexanox or vehicle control. The reduction in *Adrb3* expression (due to feedback inhibition) by amlexanox treatment was significant in both the WT and SMKO mature adipocytes, with no significant differences between the genotypes (Fig. 4A). Amlexanox induced *Il4ra* expression in WT but not SMKO ATMs (Fig. 4B). Consistent with sensitization to IL-4 signaling in vivo, amlexanox treatment increased the expression of *Il10* in ATMs from WT but not SMKO WAT (Fig. 4C). The frequency of proinflammatory Cd11c^+^ ATMs decreased 52 h after amlexanox treatment in WT but not SMKO mice, again indicating dependence on macrophage STAT3 (Fig. 4D and E). Notably, we did not observe any change in the expression of macrophage recruiting factor *Mcp1* in the adipocytes from mice treated with amlexanox versus vehicle control (Fig. 4F). These data support a model in which adipocyte-secreted IL-6 resulting from oral amlexanox treatment plays a paracrine role activating local macrophage STAT3, which in turn upregulates the expression of *Il4ra*, thereby increasing the sensitivity of the macrophages to IL-4 signaling (Fig. 4G).

**Fig. 4.**
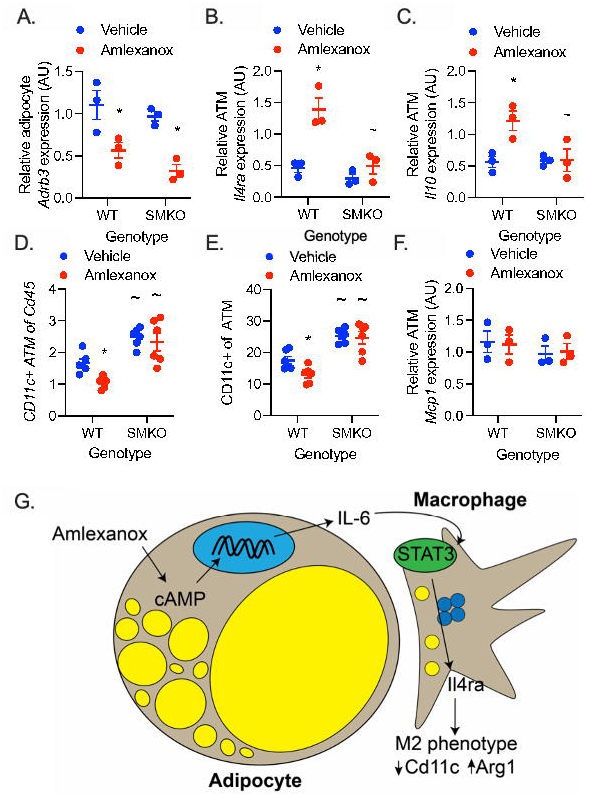
In vivo sensitization of macrophages to IL-4 by amlexanox requires STAT3. (A-C) Obese male mice aged 20-24 weeks were treated with 25 mg/kg amlexanox or vehicle control. After 4 h the eWAT was collected and digested with collagenase. n = 3 per group. (A) Q-PCR analysis of *Adrb3* expression in mature adipocytes from epididymal fat. (B-C) Q-PCR analysis of gene expression in epididymal ATMs (Cd45^+^, F4/80^+^, Cd11b^+^, Cd3^-^) isolated by FACS from SVC. (B) Relative *Il4ra* expression in ATMs. (C) Relative *Il10* expression in ATMs. (D-E) FACS analysis of SVC isolated from the epididymal fat of obese male mice aged 20-24 weeks treated with 25 mg/kg amlexanox or vehicle control for 52 hours. n = 6 per group. Macrophages defined as CD45^+^, F4/80^+^, CD11b^+^, Cd3^-^ cells. (D) CD11c^+^ macrophages as a percentage of CD45^+^ cells. (E) CD11c^+^ macrophages as a percentage of total macrophages. (F) Q-PCR analysis of *Mcp1* expression in mature adipocytes from epididymal fat. n = 3 per group. (G) Schematic of oral amlexanox treatment and activation of this adipocyte to macrophage communication axis. Statistical significance determined by post hoc analysis after significant two-way ANOVA. * p < 0.05 vehicle versus amlexanox within genotype. ^∼^ p < 0.05 WT versus KO within treatment group.

## Discussion

The role of IL-6 in adipose tissue inflammation and metabolic health is complex. Adding to this complexity, there are numerous cellular sources of IL-6 in the WAT [13, 16]. IL-6 is a pleotropic cytokine whose production is stimulated by a variety of signaling pathways. In adipocytes IL-6 secretion can be stimulated by inflammatory NF-*κ*B signaling or catabolic cAMP signaling down stream of catecholamine stimulation. Studies in obese adipocyte-specific *Il6* KO mice have suggested that adipocyte-secreted IL-6 promotes WAT inflammation or has no net effect [13, 14, 17]. Obese adipocytes primarily secrete IL-6 downstream of inflammatory not catabolic stimulation, due to increased NF-*κ*B signaling and catecholamine resistance [3, 4, 18, 19]. To probe the impact of catabolic IL-6 secretion from adipocytes in the context of obesity, we utilized amlexanox treatment which reverses intracellular catecholamine resistance to stimulate IL-6 secretion in obese adipocytes [2, 4]. We observed that amlexanox-stimulated IL-6 secretion from adipocytes activates STAT3 in ATMs, sensitizing them to IL-4 by upregulating the expression of *IL4ra*. Previous studies have reported direct anti-inflammatory effects of amlexanox [20-22], but direct amlexanox treatment in BMDMs did not activate STAT3 or sensitize them to IL-4 treatment. We did observe a direct additive effect of amlexanox to suppress *Itgax* expression in BMDMs. However, the in vivo impact of amlexanox to acutely reduce the percent CD11c^+^ ATMs was dependent on macrophage STAT3. While our results suggest an anti-inflammatory effect of amlexanox-induced IL-6 secretion from adipocytes, there are likely multiple pathways by which amlexanox affects WAT inflammation in obese animals. One limitation of these studies is the exclusive use of amlexanox to induce catabolic adipocyte IL-6 secretion. Additional studies will be required to determine whether this signaling axis is active in other physiological/pathophysiological contexts when adipocytes are catabolically activated.

## Acknowledgments

We thank the University of Michigan’s Flow Cytometry Core, the VA San Diego Healthcare System’s Flow Cytometry Research Core, UCSD’s Flow Cytometry Core in Moores Cancer Center, and the University of Michigan Cancer Center Research Histology Laboratory. We also thank Brian Zamarron, and Carey Lumeng from the University of Michigan in Ann Arbor, Michigan for assistance in designing the FACS antibody panels.

## Author Contributions

Conceptualization, S.M.R.; Formal Analysis. D.L., B.D., J.Z., P.E.B., J.H.D., J.C. and S.M.R.; Investigation D.L., B.D., J.Z., P.E.B., J.H.D., W.X., J.C. and S.M.R.; Methodology, D.L., and S.M.R.; Supervision S.M.R., Visualization S.M.R.; Writing – original draft S.M.R., B.D., J.H.D.; Writing – review & editing S.M.R., B.D., J.Z., P.E.B., and J.H.D. S.M.R. is the guarantor of this work and, as such, had full access to all the data in the study and takes responsibility for the integrity of the data and the accuracy of the data analysis.

