## Supplementary Material for "Adipocyte-secreted IL-6 sensitizes macrophages to IL-4 signaling"

### Supplementary Figures

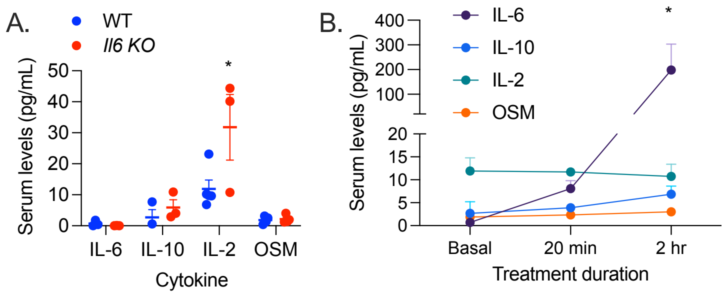


Supplemental Figure 1: **IL-6 family cytokines**. (A) Serum levels of IL-6 family cytokines in obese male WT and *Il6* KO mice. (B) Serum levels of IL-6 family cytokines in obese male mice at baseline and 20 minutes or 2 h after gavage with 25 mg/kg amlexanox. Statistical significance determined by post hoc analysis after significant ANOVA. * $p < 0.05$ WT versus *Il6* KO or baseline versus 2 h amlexanox.

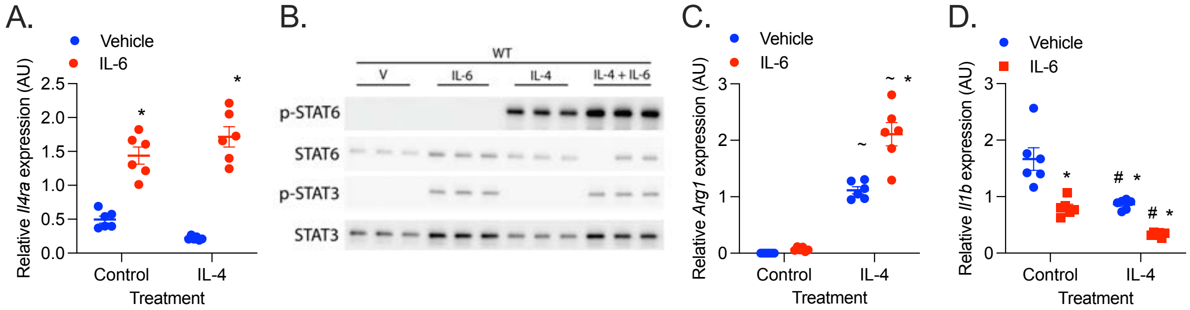


Supplemental Figure 2: **IL-6 sensitizes macrophages to IL-4**. (A) *Il4ra* expression in BMDM treated with IL-6 at day 6 for 24 h. (B) Western blot analysis of STAT3 and STAT6 phosphorylation in BMDM treated with IL-6 on day 6, then IL-4 on day 7 for 30 minutes. (C-D) Gene expression in BMDM treated with IL-6 at day 6, then IL-4 on day 7 for 24 h. (C) *Arg1* expression. (D) *Il1b* expression. Statistical significance determined by post hoc analysis after significant ANOVA. * $p < 0.05$ Vehicle versus IL-6, $^{\#}$ $p < 0.05$ Control versus IL-4.

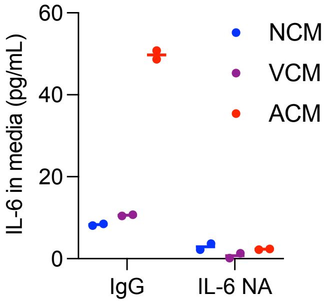


Supplemental Figure 3: **IL-6 in adipocyte-conditioned media**. Levels of IL-6 in conditioned media treated with IgG or IL-6NA. Effective neutralization is evidenced by the inability of the ELISA to detect IL-6 in the media.

**Supplemental Table 1: Q-PCR primers**

| Gene | Forward Primer | Reverse Primer |
|---|---|---|
| *Adrb3* | 5’-GGCCCTCTCTAGTTCCCAG-3’ | 5’-TAGCCATCAAACCTGTTGAGC-3’ |
| *Arbp* | 5’-CACTGGTCTAGGACCCGAGAA-3’ | 5’-AGGGGGAGATGTTCAGCATGT-3’ |
| *Arg1* | 5'-CTCCAAGCCAAAGTCCTTAGAG -3' | 5'-AGGAGCTGTCATTAGGGACATC-3' |
| *Itgax* | 5'-CTGGATAGCCTTTCTTCTGCTG-3' | 5'-GCACACTGTGTCCGAACTCA-3' |
| *Il1b* | 5'- GCAACT GTTCCTGAACTCAACT-3’ | 5'-ATC TTT TGG GGT CCG TCA ACT-3’ |
| *Il10* | 5'-CCCATTCCTCGTCACGATCTC-3' | 5'-TCAGACTGGTTTGGGATAGGTTT-3' |
| *Il4ra* | 5'-TCTGCATCCCGTTGTTTTGC-3’ | 5'-GCACCTGTGCATCCTGAATG-3’ |
| *Mcp1 (Ccl2)* | 5'-TCTCTCTTCCTCCACCACCATG-3’ | 5'-CAAGGCTCACCATCATCGTAG-3’ |
| *Socs3* | 5’-ATGGTCACCCACAGCAAGTTT-3’ | 5’-TCCACTAGAATCCGCTCTCCT-3’ |
